# The Bicaudal-D/Egalitarian complex defines the specificity of cargo transport by Dynein

**DOI:** 10.1101/2023.09.11.557182

**Authors:** Frederick C. Baker, Wen Lu, Margot Lakonishok, Hannah Neiswender, Rajalakshmi Veeranan-Karmegam, Phylicia Allen, Somayesadat Badieyan, Michael A. Cianfrocco, Vladimir Gelfand, Graydon B. Gonsalvez

## Abstract

Numerous motors of the Kinesin family contribute to plus-end microtubule transport. However, almost all transport towards the minus-end of microtubules involves a single motor, cytoplasmic Dynein (Dynein). To gain motility, Dynein must interact with activating cargo adaptors. One such adaptor is Bicaudal-D (BicD; BICD2 in humans). Mutations in BICD2 are associated with Spinal Muscular Atrophy (SMA), a degenerative motor neuron disease. BicD is autoinhibited from binding Dynein in the absence of cargo. A well-characterized cargo for BicD is the RNA binding protein, Egalitarian (Egl). Egl in conjunction with BicD links mRNA to Dynein in the *Drosophila* egg chamber and embryo. To better understand how Dynein is activated and whether BicD links additional cargo with Dynein, we defined the BicD interactome in the presence and absence of Egl. This revealed a vast number of potentially novel BicD cargos including the nucleoporin Nup358/RANBP2, a known cargo of mammalian BICD2. In strains depleted of Egl, BicD remained associated with most of its cargo including Nup358. However, the interaction of BicD with Dynein was reduced. Consequently, the localization of Nup358 and its association with Dynein was disrupted. Thus, while BicD can bind diverse cargos, linking these cargos with Dynein requires Egl. Furthermore, our studies revealed that a SMA associated mutation in the cargo binding domain of BicD enhanced the Dynein mediated localization of certain cargos but disrupted the localization of others. At the organismal level, this mutation resulted in compromised mobility. Specific transport defects might therefore underlie the etiology of BicD associated SMA.

## Introduction

The intracellular distribution of numerous mRNAs, proteins, vesicles, and organelles depend on the activity of microtubule motors. Microtubules are polarized cytoskeletal structures, and the motors that traverse them move unidirectionally towards either the plus-end or the minus-end of the microtubule. Motors of the Kinesin family are tasked with transporting cargo towards the microtubule plus end ^1,2^. The human genome encodes 45 different Kinesins, and the majority of these are plus-end motors. By contrast, transport towards the minus-end of microtubules is mostly driven by a single motor, cytoplasmic Dynein (hereafter Dynein) ^3^. The mechanism by which Dynein transports this vast diversity of cargo is an important and unresolved question.

Proteins referred to as activating cargo adaptors appear to fill this role by linking Dynein with various types of cargo ^4^. As the name suggests, these adaptors are also required for fully activating Dynein for motility. The isolated Dynein motor, a 1.4 MDa complex, has limited activity. In the absence of cargo, the motor exists in a conformation referred to as the “phi particle” ^5-7^. In this state, Dynein has minimal microtubule binding affinity. To bind and transport cargo, two additional components are required, the multi-subunit Dynactin complex, and an activating cargo adaptor ^3,8-10^. All activating cargo adaptors identified to date have a coiled coil domain that binds at the interface between Dynactin and Dynein. By doing so, cargo adaptors are thought to stabilize the trimeric complex, and to position the microtubule binding domains of Dynein for optimal motility ^4,11-14^.

One of the best studies activating adaptors is *Drosophila* Bicaudal-D (BicD) or its mammalian homolog BICD2 ^15-17^. The fly and mammalian proteins are highly homologous and are thought to function in a similar manner to activate Dynein. BicD contains three coiled-coil domains, referred to as CC1, CC2, and CC3. The N-terminal coiled-coil domain (CC1) interacts with Dynein and Dynactin, whereas the C-terminal domain (CC3) binds cargo ^10,15,18-20^ (Fig. 1A). In the absence of cargo, an intramolecular interaction between the first and third coiled-coil domains of BicD essentially blocks the Dynein binding site. Upon cargo binding, the BicD intramolecular interaction is disrupted. This effectively opens the adaptor, enabling it to interact with Dynein and Dynactin (Fig. 1B) ^20-25^.

**Figure1:**
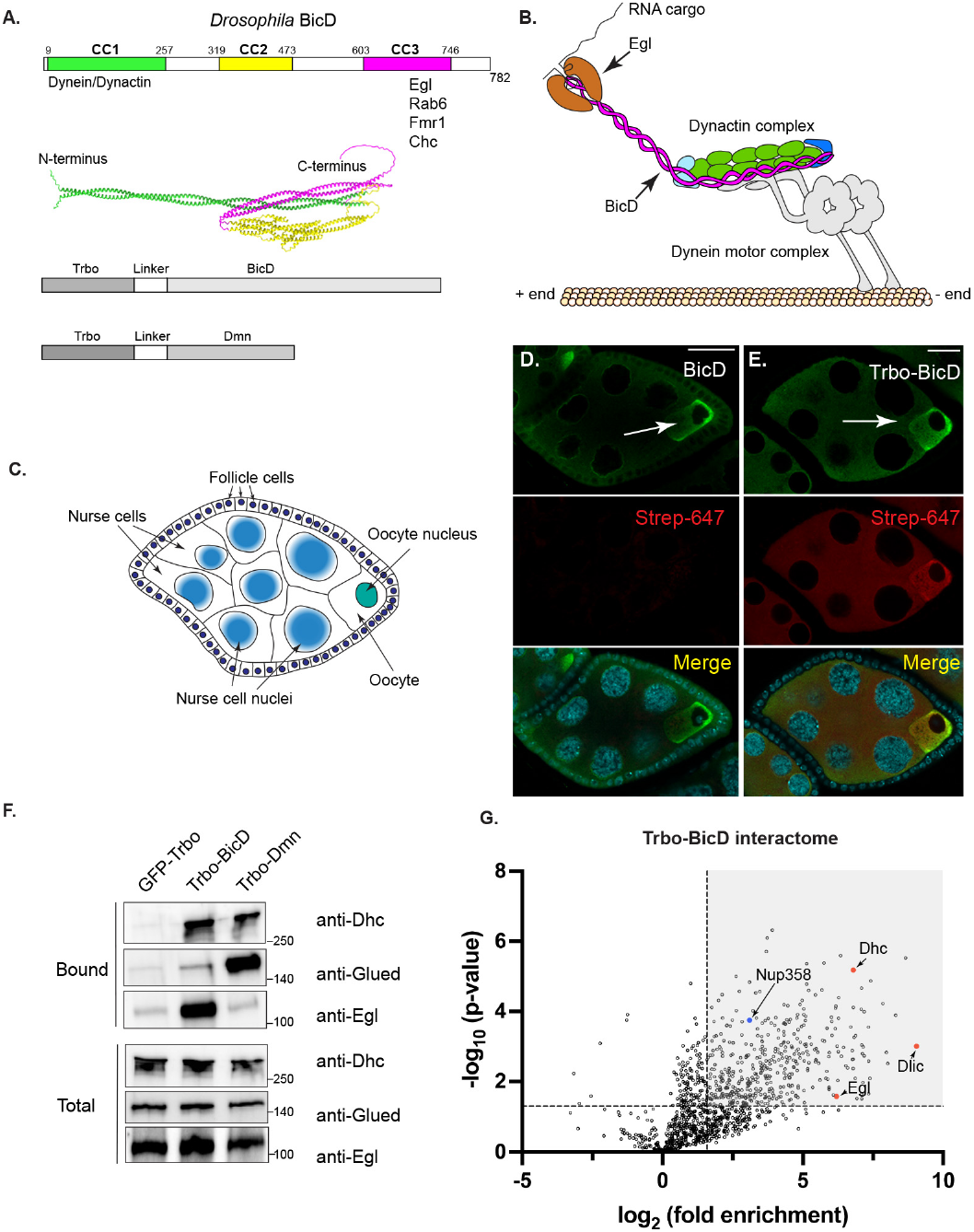
Identification of the BicD interactome. **(A)** Domain structure of *Drosophila* BicD indicating the coiled coil domains and regions bound by Dynein/Dynactin and known cargo. The middle panel depicts the structure of full-length BicD predicted using ColabFold. The model illustrates the autoinhibited conformation of BicD. The bottom panel depicts the TurboID constructs used in this work. **(B)** Schematic of the activated motor illustrating Dynein, Dynactin and cargo bound BicD. **(C)** Schematic of a *Drosophila* egg chamber. **(D)** Egg chambers from wild-type flies were processed for immunofluorescence using an antibody against BicD (green) and Streptavidin-647 (red). **(E)** Egg chambers from flies expressing Trbo-BicD were processed for immunofluorescence using a FLAG antibody (green) and Streptavidin-647 (red). The localization of Trbo-BicD is similar to endogenous BicD. **(F)** Ovarian lysates were prepared from flies expressing GFP-Trbo, Trbo-BicD or Trbo-Dmn. Biotinylated proteins were purified using Streptavidin beads and analyzed by blotting using the indicated antibodies. Total and bound fractions are shown. **(G)** Ovarian lysates were prepared from flies expressing GFP-Trbo or Trbo-BicD. The biotinylated proteins were purified as in panel F. Bound proteins were analyzed by mass spectrometry. The shaded box corresponds to proteins that have a p value of 0.05 or greater and were enriched at least three-fold with Trbo-BicD (1.585 in the log2 scale). We refer to these proteins as the BicD interactome. The scale bar is 20 microns.

The *Drosophila* ovary is an excellent model for examining Dynein based transport ^26,27^. The ovary is composed of numerous ovarioles, each of which contain egg chambers at various stages of maturation. Each egg chamber contains sixteen germline cells; fifteen nurse cells and a single oocyte. The germline cells are surrounded by a layer of somatic follicle cells (Fig. 1C) ^28^. During early stages of egg chamber maturation (stages1 through 7), the distribution of microtubules is polarized. At these stages, the minus-ends of most microtubules are localized within the oocyte ^29,30^. Thus, factors transported by Dynein at these stages are typically oocyte enriched.

Egalitarian (Egl) has been shown to link mRNA with Dynein via its interaction with BicD ^31,32^. Current models suggest that once Egl binds mRNA, it is able to associate with BicD. This relieves the BicD intramolecular interaction enabling the complex to bind Dynein and Dynactin ^23,24,33^. In addition to Egl, BicD has also been shown to interact with Rab6, Fmr1 (fragile X messenger ribonucleoprotein 1), and Clathrin heavy chain ^34-40^. BicD appears to be the primary activating cargo adaptor for Dynein in the female germline because depletion of BicD produces the same phenotype as depletion of Dynein components ^41^. Thus, it is likely that BicD links additional unknown cargo with Dynein. The goal of this work, therefore, was to determine the BicD interactome in the ovary. Furthermore, we wished to determine whether all cargo were equivalently capable of activating BicD for binding to Dynein.

## Results

### The BicD interactome in the presence and absence of Egl

To identify potential BicD cargos, we tagged BicD with TurboID on its N-terminus (Fig. 1A). TurboID is a promiscuous biotin ligase that attaches biotin modifications on lysine residues of proteins that come within 15-30nm of the tagged protein ^42^. This approach is referred to as proximity biotin ligation. We chose to attach TurboID to the N-terminus of BicD because previous studies have shown that N-terminally tagged BicD is functional and able to rescue a *bicD* null allele ^20^. Expression of our construct was driven using a maternal promoter. For simplicity, we will refer to this construct as Trbo-BicD. We also generated a strain in which Dynamitin (Dmn)/p50, a core component of the Dynactin complex, is tagged on the N-terminus with TurboID (Trbo-Dmn, Supplemental Fig. 1A, B). A strain expressing GFP-Trbo served as a control in these experiments. All constructs contained a FLAG tag for detection by blotting and immunofluorescence.

We next tested the functionality of these constructs. Ovaries were dissected from wild-type flies or flies expressing Trbo-BicD. Wild-type ovaries were processed using an antibody against endogenous BicD. Trbo-BicD ovaries were similarly processed using FLAG antibody. Both samples were also incubated with Alexa-647 conjugated Streptavidin. Streptavidin binds tightly to biotinylated protein and therefore serves as an indicator of in vivo biotinylation. Consistent with published results and with Dynein mediated transport, endogenous BicD was enriched within the oocyte (Fig. 1D) ^32^. A similar localization was detected for Trbo-BicD (Fig. 1E). Limited Streptavidin-647 signal was detected in wild-type ovaries in which no biotin ligase was expressed. However, robust Streptavidin signal was observed with Trbo-BicD (Fig1. D, E). To determine whether the tagged proteins were incorporated into endogenous complexes, biotinylated proteins were purified using Streptavidin beads. Dynein heavy chain (Dhc) and Egalitarian (Egl) were highly enriched with Trbo-BicD (Fig. 1F, Supplemental Fig. 1C, D). Glued/p150, a component of the Dynactin complex, and Dhc were highly enriched with Trbo-Dmn (Fig. 1F). Thus, Trbo-BicD and Trbo-Dmn are incorporating into endogenous motor complexes and biotinylating proximal proteins.

We next scaled up our experiment for interactome analysis. Ovaries were dissected from flies expressing GFP-Trbo or Trbo-BicD, biotinylated proteins were purified and analyzed by mass spectrometry. Proteins that were at least three-fold enriched in the Trbo-BicD pellet versus the control and had a p value of at least 0.05 were considered part of the BicD interactome (Fig. 1G, shaded box). This revealed that the BicD interactome was extensive. As expected, Dynein heavy chain (Dhc) and Dynein light intermediate chain (Dlic) were abundantly detected (Fig. 1G, Supplemental table1). Kinesin heavy chain (Khc), the motor subunit of the Kinesin-1, was also detected, consistent with results in mammalian cells (Supplemental table1) ^43^. In terms of non-motor components, Egl and Fmr1 were detected in the Trbo-BicD pellet (Fig. 1G, Supplemental table1). Nup358, a known cargo of mammalian BICD2 was also enriched in the BicD pellet (Fig. 1G, blue circle, Supplemental table1) ^43^. A GO analysis of the BicD interactome indicates an enrichment for factors with roles in “RNA localization”, “oocyte differentiation” and “microtubule based process”, consistent with known functions of BicD (Supplemental Fig. 1E) ^16,44^.

Surprisingly, Rab6 was not detected in our BicD interactome. Rab6 was one of the first cargos identified for mammalian BICD2 and is still considered to be the canonical cargo. Purified GTP-bound Rab6 was shown to bind BicD using in vitro experiments and myc-tagged Rab6 was capable of co-precipitating BicD from ovarian lysates ^20,36,37^. Our inability to detect Rab6 is not due to it being expressed at low level. In fact, at the mRNA level, Rab6 is more abundant than Egl (Supplemental Fig. 1F). Rab6 is a small protein and thus it might not have enough surface exposed lysine residues for biotinylation. To address this caveat, we used a nanobody based approach to examine the interaction between Rab6 and BicD ^45^. Ovaries were dissected from strains co-expressing GBP-Trbo and either GFP, Egl-GFP or constitutively active Rab6-EYFP. This results in tethering of GBP-Trbo to the fluorescently tagged protein and biotinylation of proximal proteins. BicD was abundantly detected with Egl-GFP but minimal binding was observed with Rab6-EYFP (Supplemental Fig. 1G). Thus, although Rab6 might be capable of interacting with BicD, it might not be a major BicD cargo in the female germline.

We recently used a similar approach to determine the Egl interactome ^45^. Many of the candidates identified in that study were also present in the BicD interactome (Supplemental Fig. 1H). To determine whether these proteins were linked to BicD via Egl, we analyzed the BicD interactome in control versus Egl depleted ovaries. Interestingly, except for a few proteins, the BicD interactome was largely unchanged (Fig. 2A). The most notable exception were components of the Dynein motor which were reduced in their association with BicD upon Egl depletion (Fig.2A, red dots). We validated this result by redoing the experiment and analyzing the pellets by western blotting. Consistent with the proteomics, the amount of Dhc and Dlic biotinylated by Trbo-BicD was reduced in the absence of Egl (Fig. 2B). By contrast, Khc was present at the same level (Fig. 2B). This suggests that in the absence of Egl, the ability of BicD to interact with Dynein (but not Kinesin) is reduced. This also suggests that many of the factors identified in the Egl interactome are most likely cargos for BicD and were purified in the earlier study because of the Egl-BicD interaction.

**Figure2:**
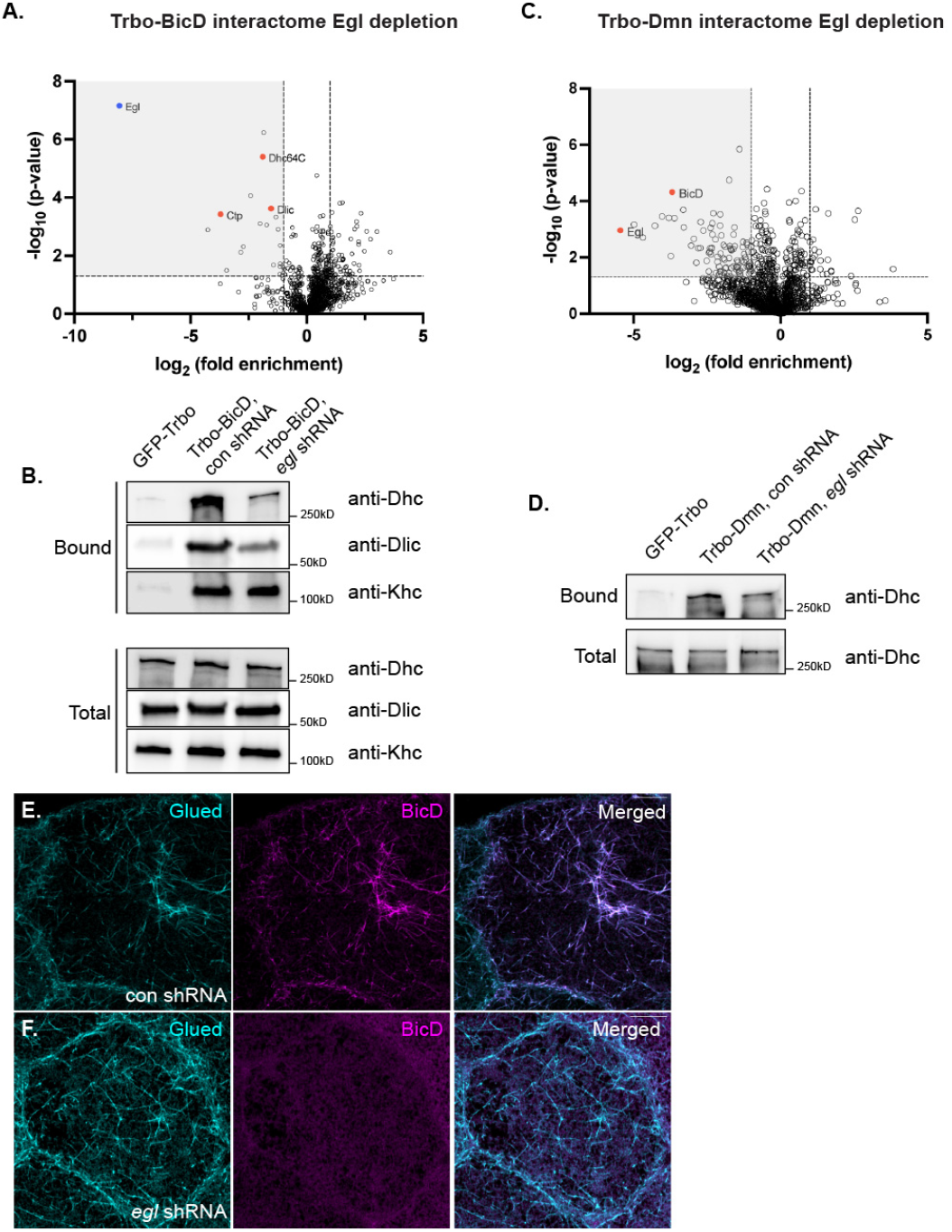
BicD and Dmn interactomes in the absence of Egl. **(A)** Ovarian lysates were prepared from flies expressing Trbo-BicD and a control shRNA or an shRNA against *egl*. Biotinylated proteins were purified using streptavidin beads and identified using mass spectrometry. **(B)** Ovarian lysates were prepared from the indicated strains, biotinylated proteins were purified and analyzed by blotting. Depletion of Egl compromises the BicD-Dynein interaction. **(C)** Ovarian lysates were prepared from flies expressing Trbo-Dmn and a control shRNA or an shRNA against *egl*. Biotinylated proteins were purified and identified as in panel A. The shaded box in panels A and C correspond to proteins that have a p value of 0.05 or greater and were reduced at least two-fold (1.0 in the log2 scale) upon Egl depletion. **(D)** Ovarian lysates were prepared from the indicated genotypes, and biotinylated proteins were purified and analyzed by blotting. The Dynein-Dynactin complex still forms upon Egl depletion. **(E-F)** Ovaries from flies expressing a control shRNA (E) or an shRNA against *egl* (F) were processed using a PIPES extraction buffer. Next the ovaries were fixed and processed for immunofluorescence using antibodies against BicD (magenta) and Glued (cyan). BicD co-localized with Glued on filaments in control but not Egl depleted egg chambers. The scale bar is 10 microns.

We next examined the Trbo-Dmn interactome in control and Egl depleted ovaries. Consistent with the above result, the level of BicD biotinylated by Trbo-Dmn was reduced in Egl depleted ovaries (Fig. 2C, Supplemental table 3). However, despite the loss of Egl, the Dynactin complex was still able to bind Dynein (Fig. 2D). It is possible that in the absence of Egl, an activating adaptor other than BicD stabilizes the Dynein-Dynactin interaction.

Treatment of egg chambers with a PIPES extraction buffer enables visualization of Egl and BicD on filamentous structures that most likely represent microtubules ^33,46^. In controls, BicD colocalized with Glued on these structures (Fig. 2E). Upon Egl depletion, Glued was still detected on filamentous structures. However, BicD was diffusely localized (Fig. 2F).

Collectively, our results suggest that despite the extensive BicD interactome, Egl is uniquely required for enabling BicD to efficiently interact with Dynein.

### Egl is not essential for bulk cytoplasmic transport

Egl is required for numerous Dynein dependent processes including oocyte specification and transport of mRNA and protein cargos into the oocyte ^30,45,47-49^. However, Dynein was recently shown to also participate in non-specific bulk cytoplasmic transport ^41^. This type of transport, which is mediated by microtubule gliding and the resulting advection forces, is required for oocyte growth. Consequently, depletion of Dhc or BicD results in a small oocyte phenotype and oogenesis arrest (Fig. 3A, B) ^41^. At early stages, Egl depletion also resulted in a small oocyte (Fig. 3C-F). However, after stage7 there was an expansion of oocyte size such that mature eggs were produced despite a loss of Egl (Fig. 3G, H). Due to the requirement of Egl in Dynein mediated mRNA localization, however, these embryos never develop ^33,47^.

**Figure3:**
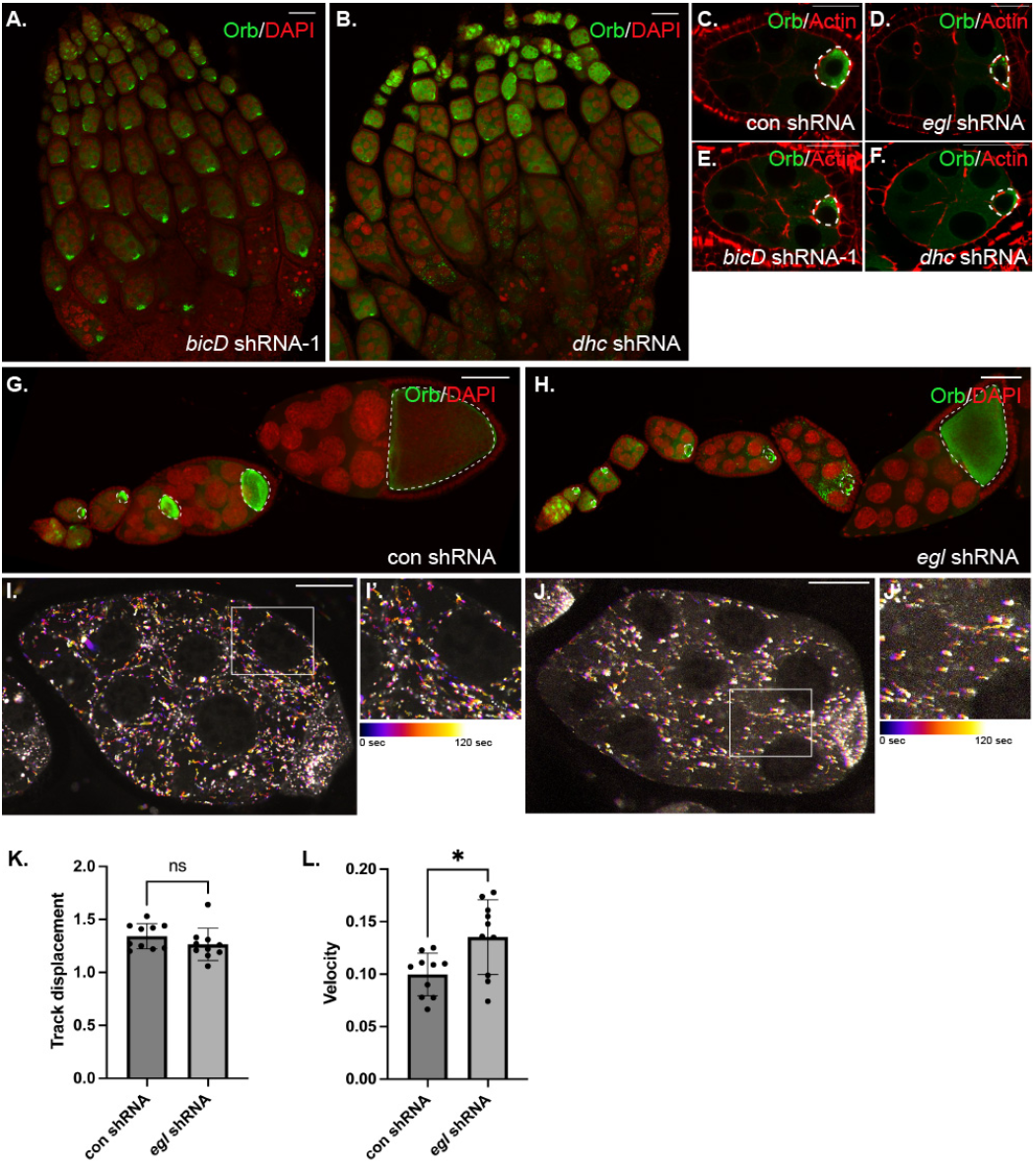
Egl is not essential for bulk cytoplasmic transport. **(A-H)** Ovaries were dissected from strains expressing a control shRNA (C, G) an shRNA against *bicD* (A, E), *dhc* (B, F), or *egl* (D, H). The ovaries were fixed and processed for immunofluorescence using Orb antibody (green). The ovaries were also counterstained using TRITC-Phalloidin which labels F-actin (red). Depletion of BicD and Dhc are associated with a small oocyte phenotype (dashed lines) and oogenesis arrest. Depletion of Egl results in a transient small oocyte phenotype but growth resumes after stage7 (dashed lines). **(I-L)** Control egg chambers (I) or those co-expressing *egl* shRNA and Sapphire tagged GEMs (J) were processed for live imaging. The panels depict the two-minute movies processed for temporal color coding. Track displacement (K) and particle velocity (L) were analyzed for both genotypes. GEMs are motile in Egl depleted egg chambers. An unpaired *t* test was used for this analysis; *p≤0.05, ns = not significant, n=10 egg chambers for each genotype. The scale bar in panels A, B, G and H is 50 microns, 10 microns in panel C-F, and 20 microns in I and J.

To examine bulk transport more directly, we visualized the movement of GEMs (Genetically Encoded Multimeric nanoparticles). GEMs are inert particles that do not directly associate with Dynein yet are moved via bulk transport ^41^. Like Dhc depletion, GEMs were less numerous in Egl depleted egg chambers (Fig. 3I, J). However, unlike Dhc depletion, in which GEM motility was virtually absent, GEM particles in Egl depleted egg chambers retained motility (Fig. 3I-K, Supplemental video 1, 2). In fact, GEMs appeared to move with a slightly higher velocity in Egl depleted egg chambers (Fig. 3L). Thus, Egl appears to be dispensable for bulk transport. Consistent with these findings, the oocyte enrichment of Golgi vesicles, a cargo likely delivered into the oocyte by bulk transport ^41^, was unaffected by Egl depletion^45^.

### Nup358 interacts with BicD and is localized in an Egl-dependent manner

Several nucleoporins were identified in the BicD interactome (Supplemental Fig. 2A, Supplemental table1). These included Nup358/RanBP2, a known cargo of mammalian BICD2. RanGAP, a direct binding partner of Nup358 ^50,51^, was also present in the BicD interactome. We verified this result by repeating the binding experiment from ovaries expressing BicD-Trbo and either a control shRNA or an shRNA targeting *egl*. As expected, Nup358 was specifically detected in the Trbo-BicD pellet. In addition, the biotinylation and precipitation of Nup358 was unchanged upon Egl depletion (Fig. 4A). Thus, depletion of Egl does not affect the BicD-Nup358 interaction.

**Figure4:**
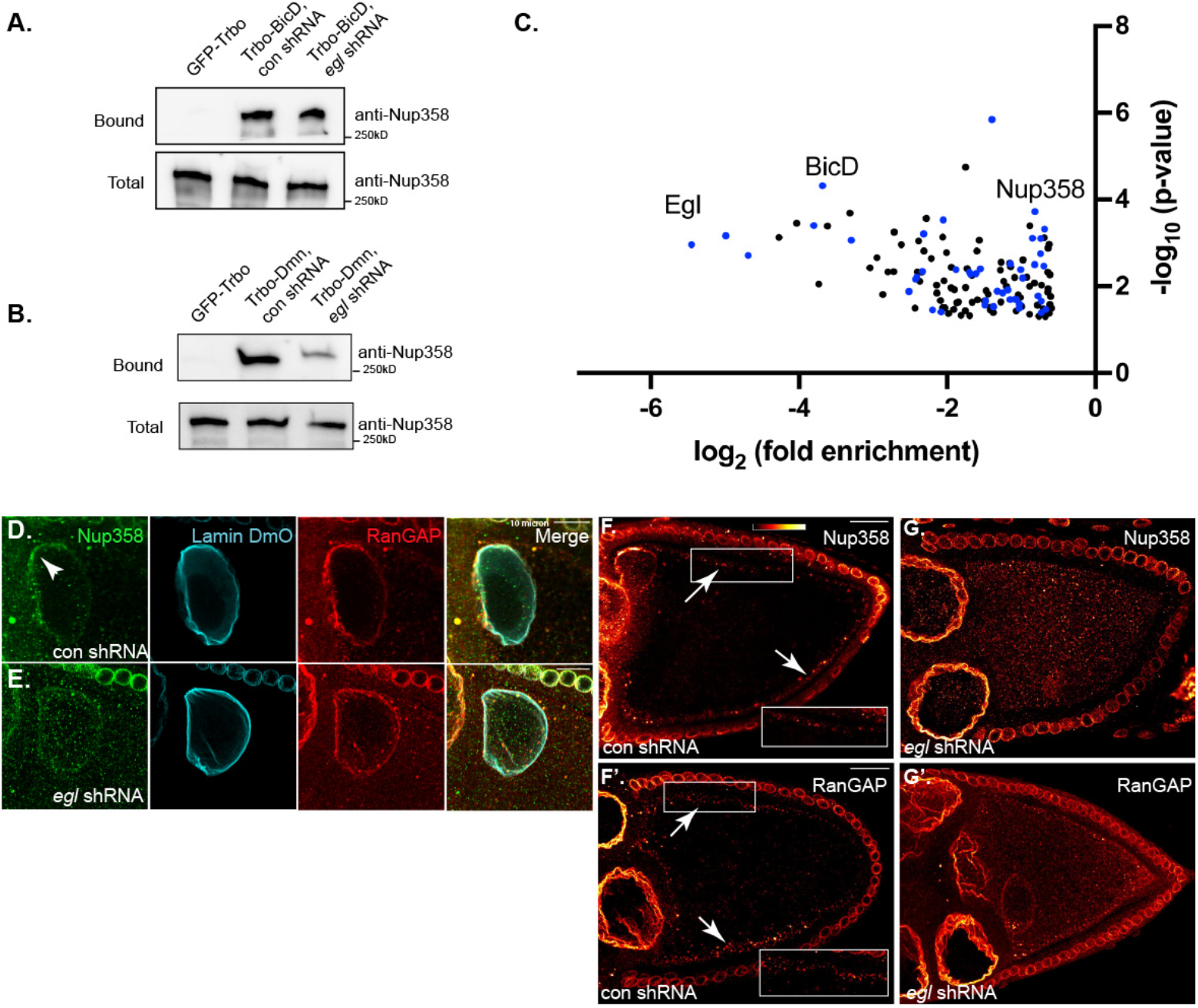
Nup358 is localized in an Egl dependent manner. **(A)** Ovarian lysates were prepared from flies expressing Trbo-BicD and either a control shRNA or an shRNA against *egl*. Biotinylated proteins were purified using streptavidin beads and were analyzed by blotting. Nup358 is biotinylated by Trbo-BicD at similar levels in control and Egl depleted strains. **(B)** Ovaries were dissected from flies expressing Trbo-Dmn and either a control shRNA or an shRNA against *egl*. The binding and western blotting was performed as in panel A. Depletion of Egl reduces the ability of Trbo-Dmn to biotinylate Nup358. **(C)** The Dmn interactome in control vs Egl depleted egg chamber initially analyzed in Fig. 2C is reshown here. In this panel, only the proteins that were significantly reduced in the Trbo-Dmn pellet in the absence of Egl are indicated. The blue dots represent proteins that are also part of the BicD interactome. **(D-E)** Ovaries from flies expressing a control shRNA (D) or an shRNA against *egl* (E) were processed for immunofluorescence using antibodies against Nup358 (green), RanGAP (red) and Lamin DmO (cyan). **(F-G)** Mid-saggital plan of stage10 egg chambers expressing a control shRNA (F, F’) or *egl* shRNA (G, G’). Signal for Nup358 and RanGAP is shown using a red to white LUT. Arrows indicate the cortical localization of Nup358 and RanGAP foci. The correct localization of Nup358 and RanGAP is dependent on Egl. The scale bar is 20 microns.

We next examined the interaction between Nup358 and Dynein. Ovarian lysates were prepared from strains co-expressing Trbo-Dmn and either a control shRNA or an shRNA targeting *egl*. In contrast to what was observed with BicD, the amount of Nup358 precipitated with Trbo-Dmn was reduced upon Egl depletion (Fig. 4B). In fact, a re-examination of the earlier Trbo-Dmn interactome revealed that several proteins that are part of the BicD interactome were reduced in their association with Trbo-Dmn in the absence of Egl (Fig. 4C, blue circles). Thus, although BicD remained bound to most of its potential cargo in the absence of Egl, the ability of the adaptor to link these cargos with Dynein was reduced.

Nup358 along with other nucleoporins were shown to localize to condensates referred to as annulate lamellae ^52^. In stage10 egg chambers, Nup358 as well as RanGAP were localized to the dorsal anterior corner of the egg chamber, around the oocyte nucleus (Fig. 4D). In the mid-saggital plane of stage10 egg chambers, Nup358 and RanGAP were localized around the oocyte cortex (Fig. 4F, F’). These localization patterns were disrupted upon Egl depletion (Fig. 4E, G, G’). Thus, although Nup358 is a BicD cargo, its correct localization in the germline requires Egl.

### Distinct germline phenotypes produced by mutants in the BicD cargo binding domain

The BicD cargo binding domain is highly conserved between flies and humans (Fig. 5A). Previous studies have shown that mutation of arginine747 in BICD2 disrupts its interaction with Nup358 ^21,53^. This residue is also noteworthy because mutations at this site are associated with spinal muscular atrophy, a degenerative neurological disorder ^54^. In *Drosophila*, this residue corresponds to arginine688 (Fig. 5A). Studies have also shown that mutation of leucine731 to alanine (L731A) in *Drosophila* BicD compromises the binding between BicD and Egl (Fig. 5A) ^20^.

**Figure5:**
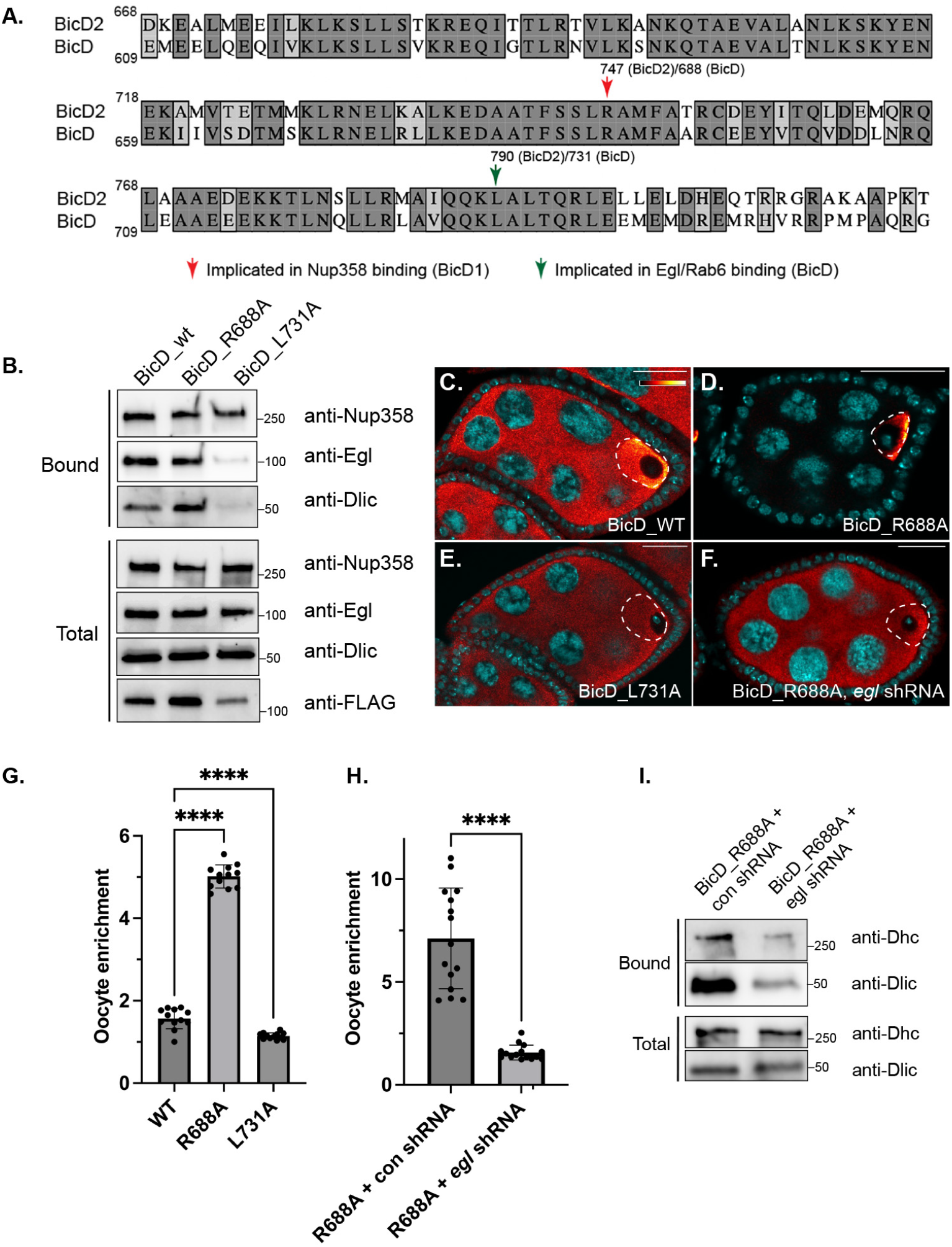
BicD cargo binding domain mutants. **(A)** Alignment of the amino acid sequences in the cargo binding domain of human BICD2 and *Drosophila* BicD. **(B)** Ovaries were dissected from flies expressing Trbo-BicD_wt, Trbo-BicD_R688A or Trbo-BicD_L731A. Lysates were prepared, biotinylated proteins were purified and analyzed by blotting using the indicated antibodies. R688A was not disrupted for interaction with Nup358 but showed increased interaction with Dlic. L731A was defective for binding to Egl and Dlic. **(C-H)** Ovaries were processed for immunofluorescence using a FLAG antibody from flies expressing Trbo-BicD_wt (C), Trbo-BicD_R688A (D), Trbo-BicD_L731A (E) or flies co-expressing Trbo-BicD_R688A and *egl* shRNA (F). The signal for FLAG is shown using a red to white LUT. The oocyte is indicated by dashed lines. Oocyte enrichment of BicD in the respective strains is quantified in panels G and H. BicD_R688A is highly enriched in the oocyte. **(I)** Ovaries were dissected from flies co-expressing Trbo-BicD_R688A and a control shRNA or an shRNA against *egl*. Lysates were prepared and biotinylated proteins were purified and analyzed by blotting using the indicated antibodies. The ability of BicD_R688A to interact with Dynein depends on Egl. The scale bar is 20 microns. A one-way Anova was used for the analysis in panel G and an unpaired *t* test was used in panel H; ****p≤0.0001, n = 12 egg chambers for panel G, n = 15 egg chambers in panel H.

To understand how these mutations affect cargo binding and Dynein activation, we generated flies expressing the respective wild-type and mutant alleles fused to N-terminal TurboID. The transgenes were expressed from the same promoter and incorporated at the same genomic locus. In contrast to what was observed in mammalian cells, mutation of R688 did not reduce the BicD-Nup358 interaction (Fig. 5B). The ability of this mutant to biotinylate Dlic was also not reduced. In fact, there was consistently more Dlic biotinylated by this mutant in comparison to wild-type (Fig. 5B). An alignment between fly and human Nup358 indicated that the proteins are poorly conserved. In fact, the specific sequence within human Nup358 required for interaction with BICD2 is not present in the *Drosophila* protein (Supplemental Fig. 2B, C) ^53^. Thus, although the interaction between BicD and Nup358 is conserved, the mechanism of interaction is not. Consistent with previous studies ^20^, mutation of L731 dramatically reduced the BicD-Egl interaction and consequently, the BicD-Dlic interaction was also affected (Fig. 5B).

As expected, wild-type BicD was enriched within the oocyte (Fig. 5C, G). In addition, as noted by Liu and colleagues ^20^, BicD_L731A was not oocyte enriched, consistent with defective Dynein binding (Fig. 5E, G). By contrast, BicD_R688A was much more highly oocyte enriched in comparison to wild-type BicD (Fig. 5D, G). In addition, a filamentous localization pattern could be detected for BicD_R688A without the need for PIPES extraction (Supplemental Fig. 3A). The phenotype of BicD_R688A is similar to the *bicd1* mutant in which phenylalanine684 is mutated to isoleucine (F684I) ^25^. Liu and colleagues found that the F684I mutant also associated with Dynein at a higher level and was more highly oocyte enriched ^20^. Thus, BicD_R688A produces a hyperactive phenotype.

We have shown in preceding sections that Egl is required for the efficient interaction between BicD and Dynein. We thus wondered whether this would also hold true for the hyperactive mutant. Depletion of Egl in the background of BicD_R688A resulted in diffuse distribution of BicD with no oocyte enrichment (Fig. 5F, H). Furthermore, the ability of BicD_R688A to biotinylate Dynein was greatly reduced upon Egl depletion (Fig. 5I). Thus, although BicD_R688A can produce a hyperactive phenotype, it still requires Egl to interact with Dynein.

Mutation at the equivalent position of R688 in humans is associated with spinal muscular atrophy ^54^. Because this mutation resides within the cargo binding domain, we wondered whether it would alter the BicD interactome. Interestingly, the interactome of BicD_R688A was largely similar to wild-type (Supplemental Fig. 3B, Supplemental table 4). Consistent with western blot results (Fig. 5B), the BicD-Nup358 interaction was unaffected and the interaction with Dlic was increased in the mutant (Supplemental Fig. 3B, Supplemental table 4). The interaction of BicD_R688A with P body components such as Me31b, Cup, Tral, and Dcp1 was also increased (Supplemental Fig. 3B, Supplemental table 4). Thus, although this mutation does not globally affect the BicD interactome, it results in increased interaction with Dynein and select cargos.

We next determined the ability of the mutants to rescue BicD depletion. The *bicD* shRNA-1 used in the preceding section produces a very strong phenotype with oogenesis arrest at early stages (Fig. 3A). This phenotype could not be rescued even with Trbo-BicD_wt (data not shown). A different shRNA targeting BicD was also available from the stock center. For simplicity, we refer to this strain as *bicD* shRNA-2. Expression of this shRNA using the same driver was capable of depleting BicD in the germline (Supplemental Fig. 4A-C). However, BicD depletion using this shRNA did not result in oogenesis arrest.

In comparison to a strain expressing a control shRNA, Egl and Dhc were much less oocyte enriched upon BicD depletion. Expression of Trbo-BicD_wt in the depletion background rescued this phenotype (Fig. 6A-C, F-H, S, Supplemental Fig. 4I). Consistent with the binding results shown in Fig. 5, Tbro-BicD_L731A failed to restore the oocyte enrichment of Dhc and Egl (Fig. 6E, J, S, Supplemental Fig. 4I). Expression of Trbo-BicD_R688A in the depletion background produced a very interesting phenotype; Egl, but not Dhc, was much more highly enriched in the oocyte (Fig. 6D, I, S, Supplemental Fig. 4I). It therefore appears that hyperactive phenotype of BicD_R688A correlates specifically to BicD cargos and not with global Dynein hyperactivation. In stage10 egg chambers, Egl could be detected on filamentous structures in nurse cells of flies expressing the control shRNA but not *bicD* shRNA-2 (Supplemental Fig. 4D, E, asterisk). The filamentous localization of Egl was restored in egg chambers co-expressing the shRNA and either Tbro-BicD_wt or Tbro-BicD_R688A, but not Tbro-BicD_L731A (Supplemental Fig. 4F-H, asterisk). Interestingly, in the hyperactive mutant, a strong accumulation of Egl was also detected at the anterior margin of the oocyte (Supplemental Fig. 4G, arrow).

**Figure6:**
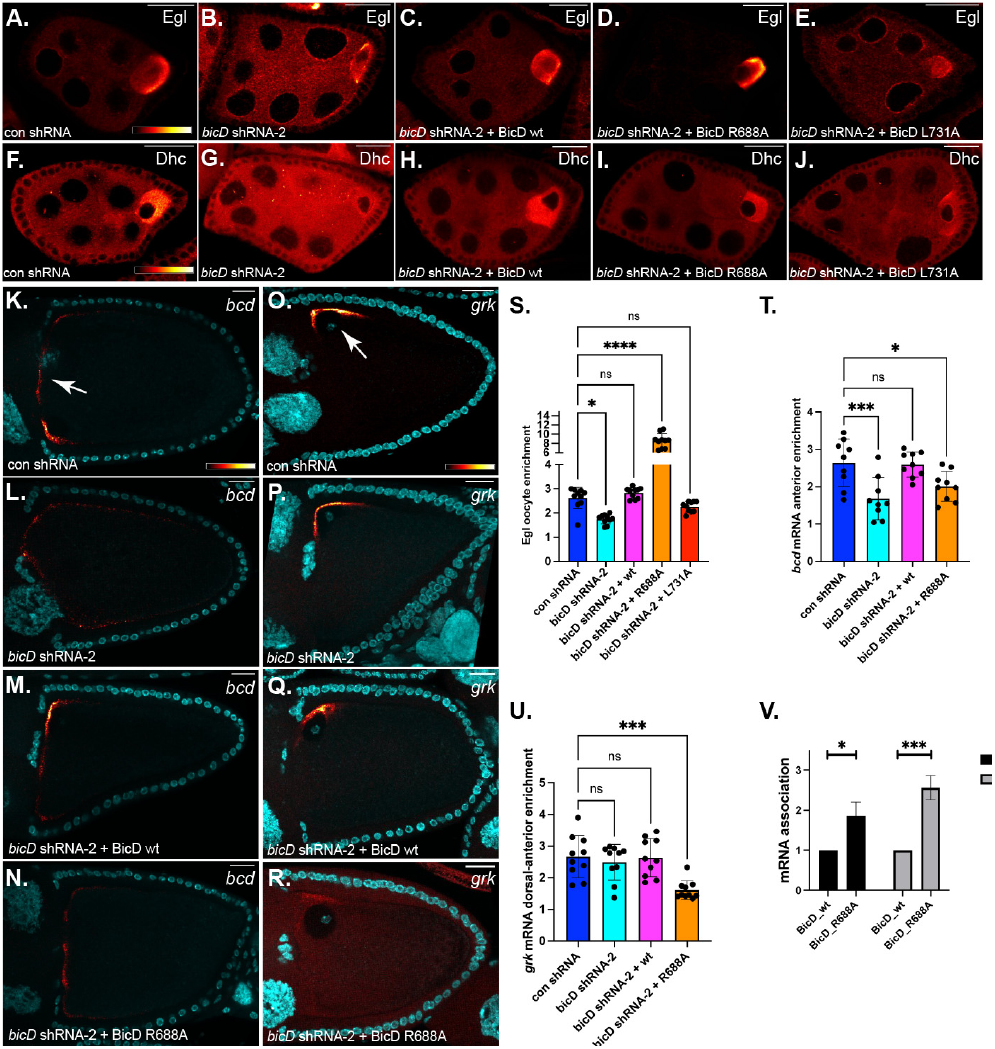
Cargo localization in BicD mutants. **(A-J)** Ovaries from strains expressing a control shRNA (A, F), an shRNA against *bicD (*B, G), or co-expressing the *bicD* shRNA along with Trbo-BicD_wt (C, H), Trbo-BicD_R688A (D, I) or Trbo-BicD_L731A (E, J), were fixed and processed using antibodies against either Egl (A-E) or Dhc (F-J). **(K-N)** Egg chambers from flies expressing a control shRNA (K), an shRNA against *bicD* (L), or co-expressing the *bicD* shRNA along with Trbo-BicD_wt (M) or Trbo-BicD_R688A (N) were fixed and processed for in situ hybridization using probes against *bicoid* mRNA (*bcd*). The arrow in panel K represents the normal localization of *bcd* mRNA at the anterior of the oocyte. **(O-R)** The same genotypes were processed using probes against *grk* mRNA. The arrow in panel O represents the normal localization of *grk* mRNA at the dorsal-anterior corner of the oocyte. **(S)** The oocyte enrichment of Egl in the indicated strains was quantified. **(T-U)** The anterior accumulation of *bcd* mRNA (T) and the dorsal-anterior enrichment of *grk* mRNA (U) was quantified in the indicated strains. Signal for Egl, Dhc, *bcd*, and *grk* mRNAs are shown using the red to white LUT. DAPI (cyan) which labels nuclei is also shown in panel K-R. **(V)** Ovaries were dissected from flies co-expressing *bicD* shRNA-2 and Trbo-BicD_wt or Trbo-BicD_R688A. Wild-type and mutant Trbo-BicD was immunoprecipitated using FLAG antibody. The co-precipitating RNA was extracted and analyzed using reverse transcription followed by qPCR. The level of *bcd* and *grk* co-precipitating with BicD-R688A was normalized to the amount co-precipitating with wild-type BicD. The entire experiment was done in triplicate (n = 3). The scale bar is 20 microns. A one-way Anova was used for the analysis in panels S, T and U and an unpaired *t* test was for used for the analysis in panel V. ****p≤0.0001, ***p≤0.001, *p≤0.05, ns = not significant, n = 10 egg chambers for panels S and U and n = 9 egg chambers for panel T.

We next examined mRNA cargos localized in an Egl/BicD dependent manner. *bicoid* (*bcd*) is localized at the anterior margin of wild-type stage10 egg chambers and those expressing a control shRNA (Fig. 6K, arrow) ^55^. Depletion of BicD results in spreading of this signal and a reduced accumulation of *bcd* at the oocyte anterior. This phenotype was rescued by Tbro-BicD_wt (Fig. 6L, M, T). Interestingly, even though Trbo-BicD_R688A interacts with Egl and Dynein, the anterior localization of *bcd* mRNA was not restored (Fig. 6N, T). *grk* is normally localized at the dorsal-anterior corner of the oocyte (Fig. 7O, arrow) ^56^. Expression of *bicD* shRNA-2 did not disrupt the localization of *grk* (Fig. 6P, U). This suggests that although BicD is depleted by this shRNA, the residual BicD present is sufficient to localize *grk* mRNA. Expression of Tbro-BicD_wt in the depletion background did not disrupt *grk* localization (Fig. 6Q, U). Unexpectedly, expression of Trbo-BicD_R688A resulted in significant delocalization of *grk* (Fig. 6R, U).

**Figure7:**
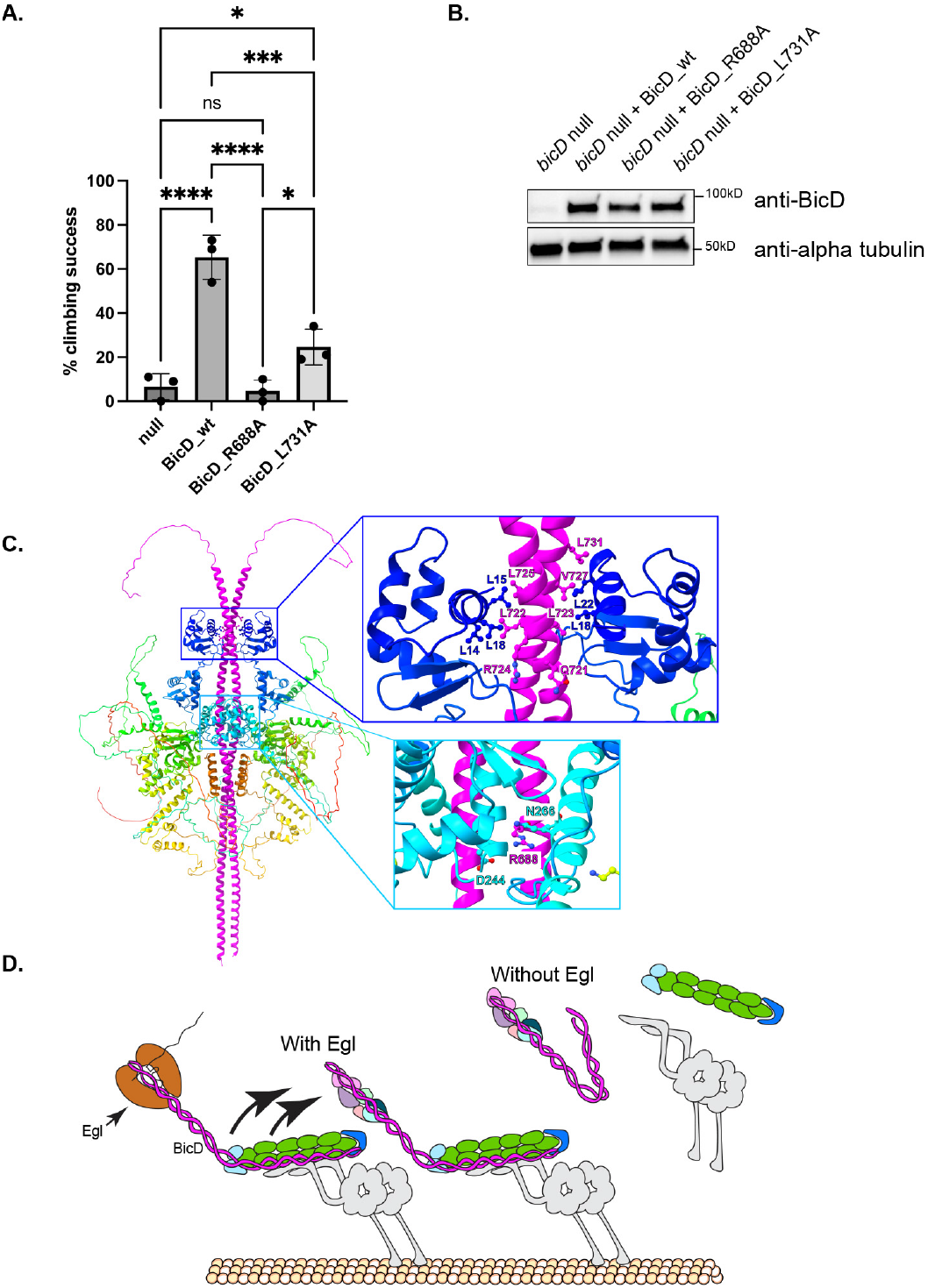
Organismal phenotype of BicD mutants. **(A)** A reverse geotaxis assay was performed on flies that were either completely null for *bicD* or expressed untagged BicD_wt, BicD_R688A, or BicD_L731A in the null background. The climbing success of each genotype is indicated. The entire experiment was performed in triplicate with each experiment including at least 9 flies per genotype. A two-way Anova was used for the analysis. ****p≤0.0001, ***p≤0.001, *p≤0.05, ns = not significant. **(B)** Lysates from these same flies were analyzed by blotting using the indicated antibodies. **(C)** ColabFold was used to predict the structure of Egl along with the cargo binding domain of BicD. BicD is represented in magenta and Egl is shown using a rainbow coloring scheme. Residues L731 and R688A on BicD are indicated along with additional residues within Egl and BicD that might participate in binding. **(D)** A model to illustrate how Egl binding to BicD might promote the recruitment of additional BicD bound cargo to the Dynein motor.

How might BicD_R688A result in mRNA delocalization? One possibility is that this mutation might cause BicD to preferentially bind to Egl that is not bound to mRNA cargo. Alternatively, the hyperactivity of this mutant might interfere with anchoring of these mRNAs. To distinguish between these possibilities, we examined the ability of wild-type and BicD_R688A to associate with these mRNAs. In comparison to wild-type, BicD_R688A was more abundantly associated with both *bcd* and *grk* mRNAs (Fig. 6V). The fact that the mRNAs are delocalized despite being bound by the Egl/BicD complex suggests that their delocalization might result from defective anchoring.

The interactome of BicD_R688A revealed that the mutant associated with P body components at a higher level in comparison to wild-type (Supplemental Fig. 3B). We therefore examined the localization of Me31b, a core P body component. In egg chambers co-expressing *bicD* shRNA-2 and Trbo-BicD_wt, Me31b was oocyte enriched in stage7 egg chambers. However, within the oocyte, Me31b was diffusely localized (Supplemental Fig. 3C) ^57^. By contrast, in stage7 egg chambers expressing *bicD* shRNA-2 and Trbo-BicD_R688A, a strong accumulation of Me31b was detected at the oocyte anterior (Supplemental Fig. 3D, arrow). A similar anterior accumulation of Me31b was also observed in stage10 egg chambers from this strain (Supplemental Fig. 3E, F, arrow). Collectively, these finds indicate that within the female germline, Egl is uniquely required for enabling BicD to interact with Dynein. In addition, a disease associated mutation in the cargo binding domain of BicD results in a hyperactive phenotype. Consequently, the localization of mRNA cargos is disrupted. However, cargos such as Egl and Me31b are excessively localized either within the oocyte in general or at the anterior margin of the oocyte.

### Organismal phenotypes of BicD mutants

BicD is expressed ubiquitously and is required for organismal health and viability ^38^. To determine the phenotypic consequences to mutations within the BicD cargo binding domain, we generated flies expressing untagged BicD_wt, BicD_R688A, or BicD_L731A. These constructs were expressed using the Dmn promoter. Dmn is a core component of the Dynactin complex. Thus, these transgenic versions of BicD should be expressed ubiquitously. These constructs were brought into the background of *bicD* nulls. As expected, *bicD* nulls that were able to emerge from the pupal case were extremely weak. A climbing assay revealed that the nulls were significantly defective in locomotor ability (Fig. 7A, Supplemental video3). By contrast, expression of a single copy of BicD_wt in the null background restored their climbing ability (Fig. 7A). Consistent with the inability of BicD_L731A to bind Dynein, expression of this mutant in the null background failed to rescue the climbing phenotype (Fig. 7A). Surprisingly, although BicD_R688A was able to interact with Dynein, expression of the hyperactive mutant in the null background resulted in flies that were unable to climb. In fact, these flies were more compromised for climbing than the L731A mutant (Fig. 7A, Supplemental video3). To determine whether lack of rescue was due to expression level differences, lysates were analyzed by western blotting. BicD_wt and BicD_L731A were expressed at similar levels.

A slight decrease was noted in the level of BicD_R688A (Fig. 7B). However, quantification of biological triplicates did not reveal a statistically significant change. Based on these findings, we conclude that the motility defect in flies expressing BicD_R688A likely results from a dominant negative effect, possibly due to Dynein mediated mis-targeting of cargo. Future studies will seek to explore the mechanistic basis of this phenotype.

## Discussion

*Drosophila* BicD and mammalian BICD2 are among the best studies Dynein activating adaptors. Research over the past decade has revealed that an intramolecular interaction within BicD keeps the protein in an inhibited conformation in the absence of cargo. Cargo binding is thought to disrupt the intramolecular interaction, enabling BicD to bind Dynein ^20-25^.

Our results suggest that although *Drosophila* BicD can bind to diverse cargos, Egl is required within the germline for enabling the BicD-Dynein interaction. Why might this be? One possibility is that the Egl-BicD interaction is abundant and other germline cargos cannot compete for binding. Consistent with this, Egl was able to efficiently co-precipitate with BicD, but Nup358 was not (data not shown). Thus, the BicD-Nup358 interaction appears to be weaker or more transient than the BicD-Egl interaction. However, if Egl was merely outcompeting other cargos, depletion of Egl should have resulted in these cargos now associating with BicD more efficiently. This was not observed, however. Our findings are therefore not compatible with a competition model.

Another possibility is that relieving BicD inhibition might be more complex than anticipated and require more than just cargo binding. For instance, the Dynein adaptor Spindly was also shown to be autoinhibited via an intramolecular interaction. Relieving this inhibition required not only binding by the RZZ complex (ROD-Zwilch-ZW10), but also association of an unknown factor present at kinetochores ^58^. Thus, a two-step activation mechanism has been proposed for Spindly ^58^. Furthermore, although mammalian BICD2 interacts directly with NUP358, the in vivo interaction of these proteins is greatly stimulated by phosphorylation events that occur during G2 phase of the cell cycle ^59^.

Our findings also suggest that in the absence of Egl, BicD remains bound to most of its cargo, but fails to link these proteins with Dynein. Thus, Egl appears to be somehow required for tethering BicD bound to distinct cargo with Dynein. This led us to wonder whether multiple cargos can be accommodated on BicD. Structural prediction using ColabFold suggests that Egl occupies a large footprint on the BicD cargo binding domain, likely making numerous contacts along the length of the protein (Fig. 7C, Supplemental, Fig. 5, Supplemental video 4) ^60^. Thus, in the presence of Egl, the same molecule of BicD likely cannot bind additional cargo. We therefore propose that formation of the Egl/BicD complex favors the binding of distinct BicD dimers with additional cargo and the linkage of these complexes with Dynein (Fig. 7D). The mechanism by which Egl accomplishes this task remains an open question. One possibility is that in the presence of Egl, BicD is post-translationally modified in a manner that promotes Dynein association. Upon Egl depletion, this modification would be lacking, and even though the adaptor remains bound to cargo, it is unable to efficiently interact with Dynein.

The function of Egl appears to be restricted to the female germline and embryo ^32,38^. *egl* nulls, although sterile, display no locomotor defect. Thus, although Egl is required for facilitating the BicD-Dynein interaction in the germline, a different mechanism is likely used in somatic tissues. Consistent with this, although we were able to detect an abundant interaction between Egl and BicD in ovaries, this was not the case in lysates from *Drosophila* heads (data not shown). Interestingly, the BicD_L731A mutant that was defective in binding to Egl and Dynein in the germline, also displayed significant locomotor defects. Another cargo might therefore bind this site in somatic tissues in an analogous manner to Egl, enabling BicD to interact with Dynein. Although we did not detect Rab6 in our germline BicD interactome, in vitro studies have shown that Rab6 and Egl bind to overlapping sites with BicD ^20^. Thus, in somatic tissues, Rab6 might be key to facilitating the BicD/Dynein interaction.

Mutation of arginine747 in humans is associated with a type of spinal muscular atrophy (SMA) ^54^. The comparable residue in *Drosophila* is arginine688 (R688), and mutation of this residue produced a hyperactive phenotype. The mutant protein was more highly enriched within the oocyte and bound Dynein at a higher level. Cargo binding, along with a hypothesized register change within the cargo binding domain of BicD, is thought to promote an open conformation, enabling the adaptor to interact with Dynein ^15,61^. R688 lies within the region hypothesized to undergo the register change ^20,62^. Thus, mutation of this residue might promote a more open BicD conformation, enabling it to interact with Dynein more efficiently. Despite this mutant displaying a hyperactive phenotype, its ability to interact with Dynein and to localize within the oocyte still required Egl. At the organismal level, flies bearing this mutation were weak and were as compromised for climbing as *bicD* nulls. Because this mutant can interact with Dynein, locomotor defects likely arise due to a dominant negative effect. Although our study did not address the somatic BicD interactome, within the germline, BicD_R688A interacted with P body components at a higher level. This resulted in inappropriate accumulation of Me31b at the oocyte anterior. P bodies are sites of translational regulation ^63^. Thus, mis-targeting P bodies within motor neurons and associated translational changes might underlie the etiology of BicD-associated SMA.

## Acknowledgements

We would like to thank Drs. Jordan Raff, Alexei Arnaoutov, and Marry Dasso for providing antibodies against Dlic, Nup358 and RanGAP. We are also grateful to the Bloomington Stock Center and the Developmental Studies Hybridoma Bank for providing fly strains and antibodies. This work was supported by grants from the National Institutes of Health to G.B.G (R35GM145340), and V.I.G. (R35GM131752). This paper was typeset with the bioRxiv word template by @Chrelli: www.github.com/chrelli/bioRxiv-word-template

## Materials and Methods

### DNA constructs

Trbo constructs were cloned into the pAttB vector ^64^. A fragment containing the promoter and 3’UTR for alpha-Tubulin67c as well as the SV40 poly adenylation sequence was generated by gene synthesis and cloned into the pAttB vector using Gibson assembly (NEB). Sequences corresponding to the cDNA for either GFP, wild-type BicD, or Dmn were then cloned using PCR and subsequent Gibson assembly in between the promoter and 3’UTR of alpha-Tubulin67c. Next, the TurboID sequence, codon optimized for expression in *Drosophila*, was generated by gene synthesis and cloned into the above constructs using Gibson assembly. This generated maternally driven GFP-Trbo, maternally driven wild-type Trbo-BicD, and maternally driven Trbo-Dmn. The R688A and L731A mutants in BicD were generated using the above wild-type Trbo-BicD construct as a template for PCR. Small fragments containing the desired mutations were generated using gene synthesis and ligated into the PCR product using Gibson assembly. This generated maternally driven Trbo-BicD_R688A and Trbo-BicD_L731A. The constructs for constitutive expression of BicD_wt and mutants were generated using a custom made pAttB vector. A region corresponding to the promoter of *dmn* as well as the 5’UTR and 1^st^ intron along with the SV40 3’UTR and polyadenylation sequence was generated by gene synthesis. Kpn1 and Not1 restriction sites were designed into this construct in between the *dmn* 5’UTR and the SV40 3’UTR. The coding sequences for BicD_wt, BicD_R688A and BicD_L731A were cloned into these restriction sites. All gene synthesized fragments were generated by Genewiz/Azenta. All constructs were verified by sequencing prior to generation of transgenic flies. The pUASp-GFP-Trbo strain was described previously ^45^.

### Fly stocks

Oregon-R-P2 (Bloomington stock center; #2376) was used as the wild-type control. Fly crosses were maintained at 25^0^C. For examination of ovarian phenotypes, female flies were dissected at 3 to 4 days of age. The pAttB-Mat-GFP-Trbo, pAttB-Mat-Trbo-BicD_WT, and pAttB-Mat-Trbo-Dmn constructs were integrated at site ZH-68E (Bloomington stock center; #24485) to generate the respective stocks. In addition, pAttB-Mat-Trbo-BicD_WT, pAttB-Mat-Trbo-BicD_R688A, and pAttB-Mat-Tbro-BicD_L731A constructs were integrated at site 86Fb (Bloomington stock center; # 24749). The transgenic strains were generated by BestGene Inc. The following shRNA strains were used:

*egl* shNRA (Bloomington stock center; #43550, donor TRiP)

*bicD* shRNA-1 (Bloomington stock center; #35405, donor TRiP)

*bicD* shRNA-2 (Bloomington stock center; # 42929, donor TRiP)

*dhc* shRNA (Bloomington stock center; #36583, donor TRiP)

*eb1* shRNA (Bloomington stock center; #36680, donor TRiP)

The *eb1* shRNA strain was used as the control strain. The strains for expression of constitutively active YFP-Rab6 (#9776) and endogenously tagged GFP-Me31b (#51530) were obtained from the Bloomington stock center. The pUASp-GFP, pUASp-Egl-GFP, pUASp-GBP-Trbo, and pUASp-Sapphire-GEM strains were previously described ^41,45,65^. Expression of UASp constructs as well as the shRNA constructs were driven using *mat atub-Gal4[V37]* (Bloomington Stock Center, #7063). For live imaging of GEMs, young adult females of the following genotypes were used; (1) *yw; UASp-F-Tractin-tdTomato/+; mat atub-Gal4[V37]/UASp-GEM (M1)* (controls), and (2) *yw; UASp-F-Tractin-tdTomato/+; mat atub-Gal4[V37]/UASp-GEM (M1), egl-shRNA* (Egl depleted). *bicD* nulls were obtained by crossing *bicD*^*r5*^/CyO (Bloomington stock center; #4553)flies to the deficiency strain, Df(2L)Exel7068/CyO (Bloomington stock center; #7838).

### Antibodies

The following antibodies were used: mouse anti-BicD 1B11 (Developmental studies hybridoma bank, 1:30 for immunofluorescence), mouse anti-BicD 4C2 (Developmental studies hybridoma bank, 1:30 for immunofluorescence), mouse anti-Dhc 2C11-2 (Developmental studies hybridoma bank, 1:300 for immunofluorescence, 1:3000 for western), mouse anti-Orb (Developmental studies hybridoma bank, 1:30 for immunofluorescence), mouse anti-LaminDmO ADL84.12 (Developmental studies hybridoma bank, 1:100 for immunofluorescence), mouse anti-GFP (Clontech,1:6,000), mouse anti-FLAG (Millipore-Sigma, 1:500 for immunofluorescence, 1:5000 for western), rabbit anti-Egl (generated inhouse, 1:500 for immunofluorescence, 1:5000 for western), rabbit anti-Khc (1:5,000 for western) ^66^, rabbit anti-Glued (1:5000 for western) ^67^, rabbit anti-Dlic (1:5000 for western, gift from Dr. Jordan Raff), rabbit anti-RanGAP (1:5000 for western, gift from Drs. Alexei Arnaoutov, and Marry Dasso), chicken anti-Nup358 (1:5000 for western, gift from Drs. Alexei Arnaoutov, and Marry Dasso). The following secondary antibodies were used: goat anti-rabbit Alexa 488, 555, and 594, (Life Technologies, 1:400); goat anti-mouse Alexa 488, 555, and 594 (Life Technologies, 1:400); goat anti-Chicken 488 (Life Technologies, 1:400); goat anti-mouse HRP (Pierce, 1:5000); goat anti-rabbit HRP (Pierce, 1:5000), and goat anti-chicken HRP (Pierce, 1:5000). Biotinylated proteins were detected in localization experiments using Streptavidin-Alexa674 (Life Technologies, 1:1200).

### Purification and analysis of biotinylated proteins

Biotinylated proteins were purified from ovarian lysates as described previously (ref). In brief, lysates were prepared using RIPA buffer (50 mM Tris-Cl [pH 7.5], 150 mM NaCl, 1% NP-40, 1 mM EDTA) containing a Halt Protease inhibitor cocktail (Piece). For small scale purification, typically about 1mg of ovarian lysate was used. The lysate was incubated with 15ul of High-Capacity Streptavidin Agarose beads (Piece) in RIPA buffer. The binding was performed overnight at 4^0^C. The samples were washed four times using RIPA buffer, bound proteins were eluted in Laemmli buffer and analyzed by western blotting. All western blot images were acquired on a Bio Rad ChemiDoc MP.

For proteomics experiments, approximately 100 ovaries were dissected for each genotype. Five mg of ovarian lysates were used in each binding experiment. Biotionylated proteins were purified using 50ul of High-Capacity Streptavidin Agarose beads, incubated at 4^0^C overnight. The following day, the samples were extensive washed using 1ml of the following: 3 times with RIPA buffer, 3 times with 1% SDS, 3 times with RIPA buffer, 3 times with high salt RIPA buffer (50 mM Tris-Cl [pH 7.5], 1M NaCl, 1% NP-40, 1 mM EDTA), 2 times with RIPA buffer and 4 times with PBS. The entire experiment was done in triplicate for proteomic analysis.

### Mass spectrometry

The mass spectrometry was performed at the Emory Integrated Proteomics Core.

#### On bead digestion

A published protocol was followed for on bead digestion of proteins ^68^. A digestion buffer containing 50 mM NH_4_HCO_3_ was added to the beads. The mixture was then with 1 mM dithiothreitol (DTT) at room temperature for 30 minutes, followed by addition of 5 mM iodoacetimide (IAA). This mixture was incubated at room temperature for an additional 30 minutes in the dark. Proteins were digested with 1 μg of lysyl endopeptidase (Wako) at room temperature overnight and were further digested overnight at room temperature with 1 μg trypsin (Promega). The resulting peptides were desalted with an HLB column (Waters) and were dried under vacuum.

#### LC-MS/MS

The data acquisition by LC-MS/MS was adapted from a published procedure ^69^. Derived peptides were resuspended in the loading buffer (0.1% trifluoroacetic acid, TFA) and were separated on a Water’s Charged Surface Hybrid (CSH) column (150 μm internal diameter (ID) x 15 cm; particle size: 1.7 μm). The samples were run on an EVOSEP liquid chromatography system using the 15 samples per day preset gradient and were monitored on a Q-Exactive Plus Hybrid Quadrupole-Orbitrap Mass Spectrometer (ThermoFisher Scientific). The mass spectrometer cycle was programmed to collect one full MS scan followed by 20 data dependent MS/MS scans. The MS scans (400-1600 m/z range, 3 × 10^6^ AGC target, 100 ms maximum ion time) were collected at a resolution of 70,000 at m/z 200 in profile mode. The HCD MS/MS spectra (1.6 m/z isolation width, 28% collision energy, 1 × 10^5^ AGC target, 100 ms maximum ion time) were acquired at a resolution of 17,500 at m/z 200. Dynamic exclusion was set to exclude previously sequenced precursor ions for 30 seconds. Precursor ions with +1, and +7, +8 or higher charge states were excluded from sequencing.

#### MaxQuant

Label-free quantification analysis was adapted from a published procedure (Seyfried, Dammer et al. 2017). Spectra were searched using the search engine Andromeda, integrated into MaxQuant, against 2020 *Drosophila melanogaster* (Fruit fly) Uniprot database (42,676 target sequences). Methionine oxidation (+15.9949 Da), asparagine and glutamine deamidation (+0.9840 Da), and protein N-terminal acetylation (+42.0106 Da) were variable modifications (up to 5 allowed per peptide); cysteine was assigned as a fixed carbamidomethyl modification (+57.0215 Da). Only fully tryptic peptides were considered with up to 2 missed cleavages in the database search. A precursor mass tolerance of ±20 ppm was applied prior to mass accuracy calibration and ±4.5 ppm after internal MaxQuant calibration. Other search settings included a maximum peptide mass of 6,000 Da, a minimum peptide length of 6 residues, 0.05 Da tolerance for orbitrap and 0.6 Da tolerance for ion trap MS/MS scans. The false discovery rate (FDR) for peptide spectral matches, proteins, and site decoy fraction were all set to 1 percent.

#### Quantification settings were as follows

re-quantify with a second peak finding attempt after protein identification has completed; match MS1 peaks between runs; a 0.7 min retention time match window was used after an alignment function was found with a 20-minute RT search space. Quantitation of proteins was performed using summed peptide intensities given by MaxQuant. The quantitation method only considered razor plus unique peptides for protein level quantitation.

### Immunofluorescence and in situ hybridization

Immunofluorescence and single molecule fluorescent in situ hybridizations (smFISH) were performed as described previously ^33^. Stellaris smFISH probes against *bcd, grk* and *osk* mRNA were obtained from LGC biosearch technologies.

### Microscopy

Fixed images were captured on either an inverted Leica Stellaris confocal microscope or an inverted Zeiss LSM780 equipped with Airyscan housed in the Augusta University Cell Imaging Core. For live images of GEMs, young adult females of the indicated genotypes (see fly stocks section) were mated with several male flies and fed active dry yeast for 16∼18 hours at room temperature (23∼24 °C) before dissection. The ovaries were dissected in Halocarbon oil 700 (Sigma-Aldrich, Cat# H8898) as previously described ^70,71^. Freshly dissected samples were imaged on a Nikon W1 spinning disk confocal microscope (Yokogawa CSU-X1 with pinhole size 50 μm) with a Hamamatsu ORCA-Fusion Digital CMOS Camera, and a 40X 1.25 N.A. silicone oil lens, with one frame every 2 seconds for 2 min, controlled by Nikon Elements software. Images were processed for presentation using Fiji, Adobe Photoshop, and Adobe Illustrator.

### RT-qPCR

Total ovarian RNA was isolated from wild-type flies using Trizol (ThermoFisher) according to the instruction provided by the manufacturer. The RNA was reverse transcribed using random hexamers and Superscript III (ThermoFisher) according to directions provided by the manufacturer. Quantitative (qPCR) was performed using SsoAdvanced Universal SYBR Green Supermix (Bio-Rad) using a Bio-Rad CFX96 Real-Time PCR. The expression level of Egl, Rab6 and Fmf1 was determined by comparing ct values for each RNA to that obtained for γ-tubulin.

### Quantifications

Enrichment of Egl, BicD, and Dhc within the oocyte was quantified by measuring the average pixel intensity of the oocyte-localized signal and dividing by the average pixel intensity of the signal in the remainder of the egg chamber. The anterior enrichment of *bcd* mRNA was determined by measuring the average pixel intensity at the anterior margin of the oocyte (localized signal) divided by the average pixel intensity within the remainder of the oocyte (delocalized signal). The dorsal-anterior enrichment of *grk* mRNA was similarly quantified by measuring the average pixel intensity at the dorsal anterior region of the oocyte (localized signal) divided by the average pixel intensity within the remainder of the oocyte (delocalized signal). ImageJ/Fiji was used for this analysis. Graphpad Prism9 was used for statistical analysis as well as generation of graphs. Track displacement and track velocity for GEMs was determined using the Imaris10 software (Oxford instruments). The “spots” module in Imaris was used to define and track GEM particles.

### *Drosophila* negative geotaxis assay

To assess adult fly motility we used a Rapid Iterative Negative Geotaxis (RING) protocol modified from ^72^. Ten newly emerged adult flies (5 males and 5 females flies) of the appropriate genotype were collected under light CO_2_ anesthetization and placed in a polystyrene vial containing our standard fly food. Flies were maintained for 3 days at 25^0^C to allow for recovery from CO_2_. Flies were then transferred without anesthetizing to newly prepared polystyrene vials without food. The vials were then assembled into a 3D printed apparatus with the vial opening oriented to the bottom of the apparatus and sealed with a cotton plug. Following a 30-minute acclimation period, the apparatus was sharply tapped on the bench surface 3 times to knock the flies to the bottom of the vial. The flies were then scored based on the percentage of flies that were able to successfully climb 50% of the length of the vial in 10 seconds. This value was averaged over 7 trials each performed 1 minute apart. The entire experiment was repeated in triplicate (n=3).

### Structure prediction

The structure of full-length BicD shown in Fig. 1A and the BicD-Egl interaction shown in Fig.7C were modeled using the ColabFold tool ^60^ integrated in the Cosmic2 web platform ^73^. Only the CC3 domain of BicD was used for the analysis in Fig.7C. Diagnostic plots (sequence coverage, alignment error, and per residue confidence score) for evaluation of the predicted data is presented in Supplemental Fig. 5.

