## Supplemental figure legends for "The Bicaudal-D/Egalitarian complex defines the specificity of cargo transport by Dynein"

**Supplemental Figure1:**
**(A-B)** Ovaries from flies expressing either GFP-Dmn (A) or Trbo-Dmn (B) were fixed and processed. The GFP tagged flies were processed using an antibody against. The Trbo tagged flies were processed using the FLAG antibody. Both constructs localize in a similar manner. **(C-D)** Ovaries were dissected from flies expressing either a control shRNA (C) or an shRNA against *egl* (D). The ovaries were fixed and processed using the homemade Egl antibody. Signal for Egl is greatly reduced in the germline of flies expressing the *egl* shRNA. **(E)** A "biological function" GO analysis was performed on the BicD interactome. The top 10 enriched categories are shown. **(F)** RNA was extracted from wild-type flies, reverse transcribed, and analyzed using quantitative PCR. The entire experiment was done in triplicate. The level of Rab6 and Egl were normalized to the level of Gamma tubulin mRNA. **(G)** Ovarian lysates were prepared from the indicated genotypes. Biotinylated proteins were purified and analyzed by western blot using the indicated antibodies. **(H)** A pie chart illustrating the proteins found as part of either the BicD interactome (this work) or the Egl interactome [45]. The scale bar is 20 microns. An unpaired *t* test was used for this analysis; ****$p \leq 0.0001$.

**Supplemental Figure2:**
**(A)** Schematic of the Nucleoporins identified in the BicD interactome. A STRING analysis was performed on these proteins to reveal known interactions. **(B)** Schematic of the domain organization of human and fly Nup358. **(C)** An alignment analysis between human and fly Nup358. The specific sequences within Nup358 that were shown to interact with BicD2 and Klc are indicated.

**Supplemental Figure3:**
**(A)** BicD_R688A localizes to filaments in the nurse cell cytoplasm. The localization pattern is seen without PIPES extraction. **(B)** Ovaries were dissected from flies expressing Trbo-BicD_wt or Trbo-BicD_R688A. Biotinylated proteins were purified and identified using mass spectrometry. **(C-F)** Localization of GFP-Me31b in strains co-expressing *bicD* shRNA-2 and Trbo-BicD_wt (C, E) or Trbo_BicD_R688A (D, F). The arrow indicates the anterior accumulation of Me31b. The scale bar is either 10 microns (A) or 20 microns (C-F).

**Supplemental Figure4:**
**(A-C)** Ovaries were dissected from flies expressing a control shRNA or *bicD* shRNA-2. One set was processed for western blot using the indicated antibodies (A). The other set was processed for immunofluorescence using the BicD antibody (B, C). **(D-H)** Ovaries from flies expressing a control shRNA (D), an shRNA against *bicD* (E), or co-expressing the *bicD* shRNA along with Trbo-BicD_wt (F), Trbo-BicD_R688A (G) or Trbo-BicD_L731A (H), were fixed and processed using an antibody against Egl. The arrow indicates the anterior oocyte localization of Egl. Signal for BicD and Egl are shown using the red to white LUT. **(I)** Quantification of Dhc oocyte enrichment from the panels shown in Fig. 6F-J. A one-way Anova was used for this analysis. ****$p \leq 0.0001$, *$p \leq 0.05$, ns = not significant, n = 10 egg chambers each genotype.

**Supplemental Figure5:**
Diagnostic plots for sequence coverage (A), alignment error (B), and per residue confidence score (C) are presented for the ColabFold model shown in Fig.7C.

**Supplemental Table1:**
The BicD interactome.

**Supplemental Table2:**
The BicD interactome in ovaries from control flies versus flies depleted of Egl.

**Supplemental Table3:**
The Dmn interactome in ovaries from control flies versus flies depleted of Egl.

**Supplemental Table4:**
The interactome of BicD_wt vs BicD_R688A.

**Supplemental Video1:**
GEM movement in control egg chambers.

**Supplemental Video2:**
GEM movement in Egl depleted egg chambers.

**Supplemental Video3:**
Negative geotaxis assay using the following genotypes; *bicD* nulls (vial 2), *bicD* nulls + BicD_wt (vial 1), *bicD* nulls + BicD_R688A (vial 3), and *bicD* nulls + BicD_L731A (vial 4).

**Supplemental Video4:**
ColabFold model of the BicD-Egl interaction.
