## Supplemental figures for "The Bicaudal-D/Egalitarian complex defines the specificity of cargo transport by Dynein"

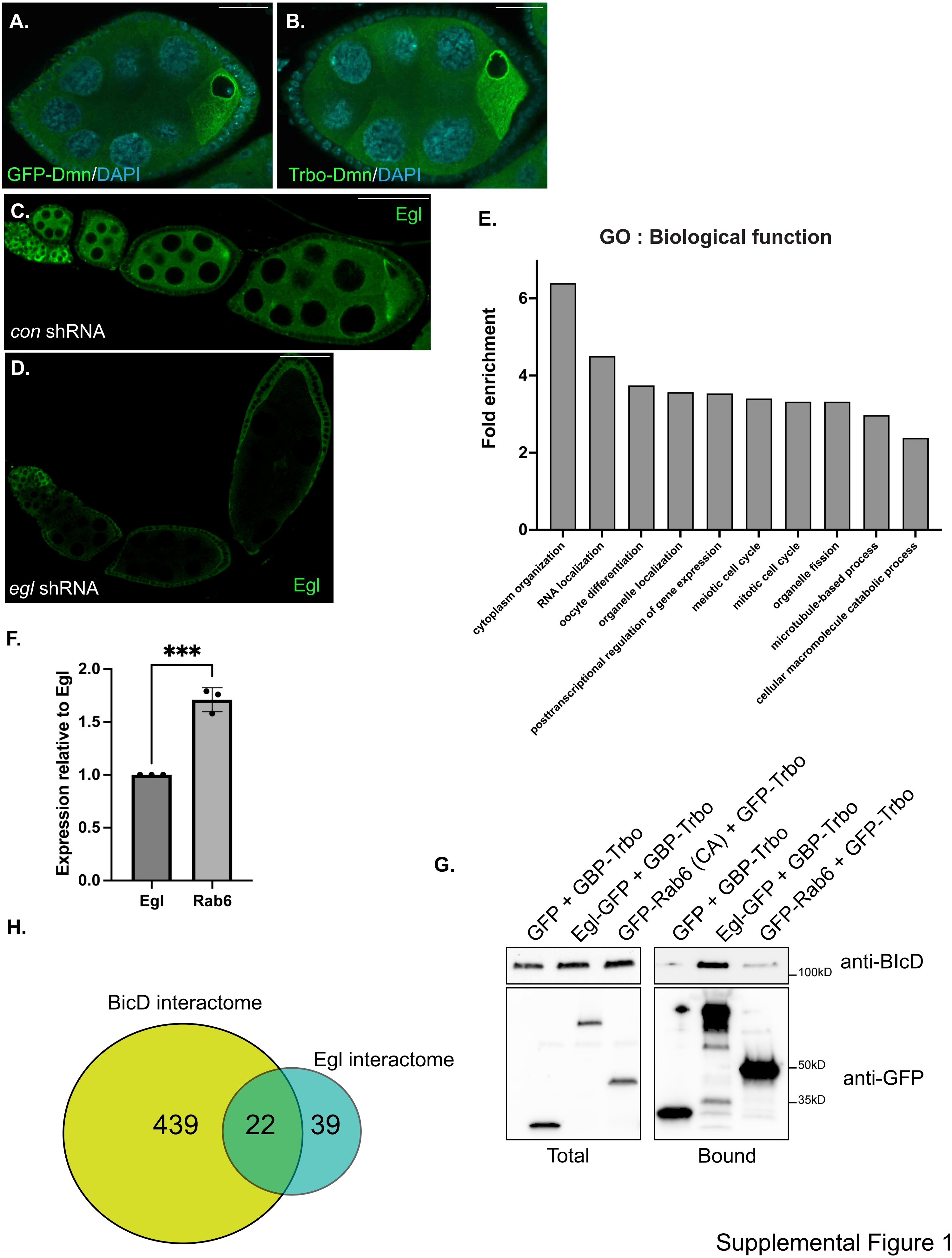


Supplemental Figure 1

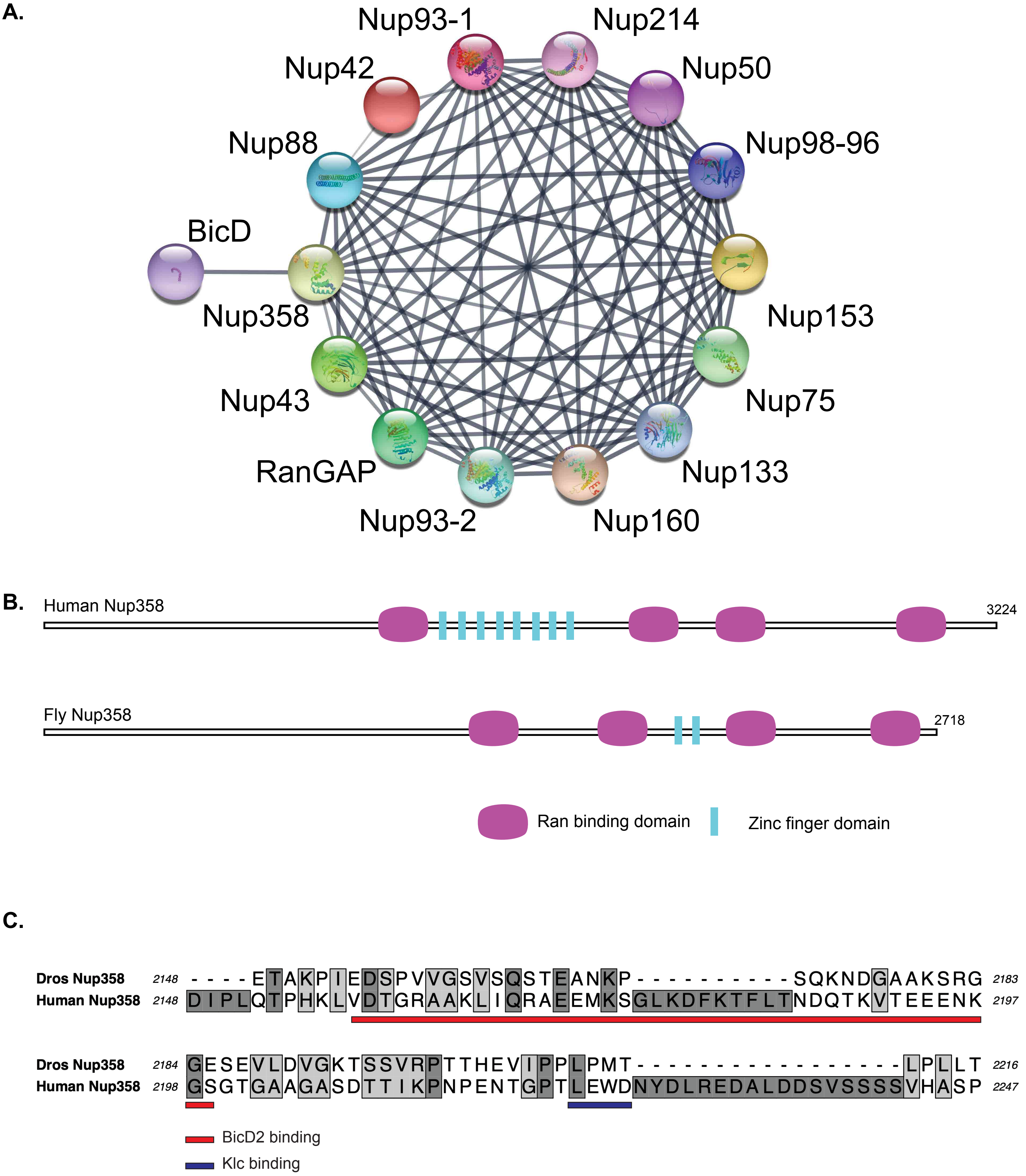


Supplemental Figure 2

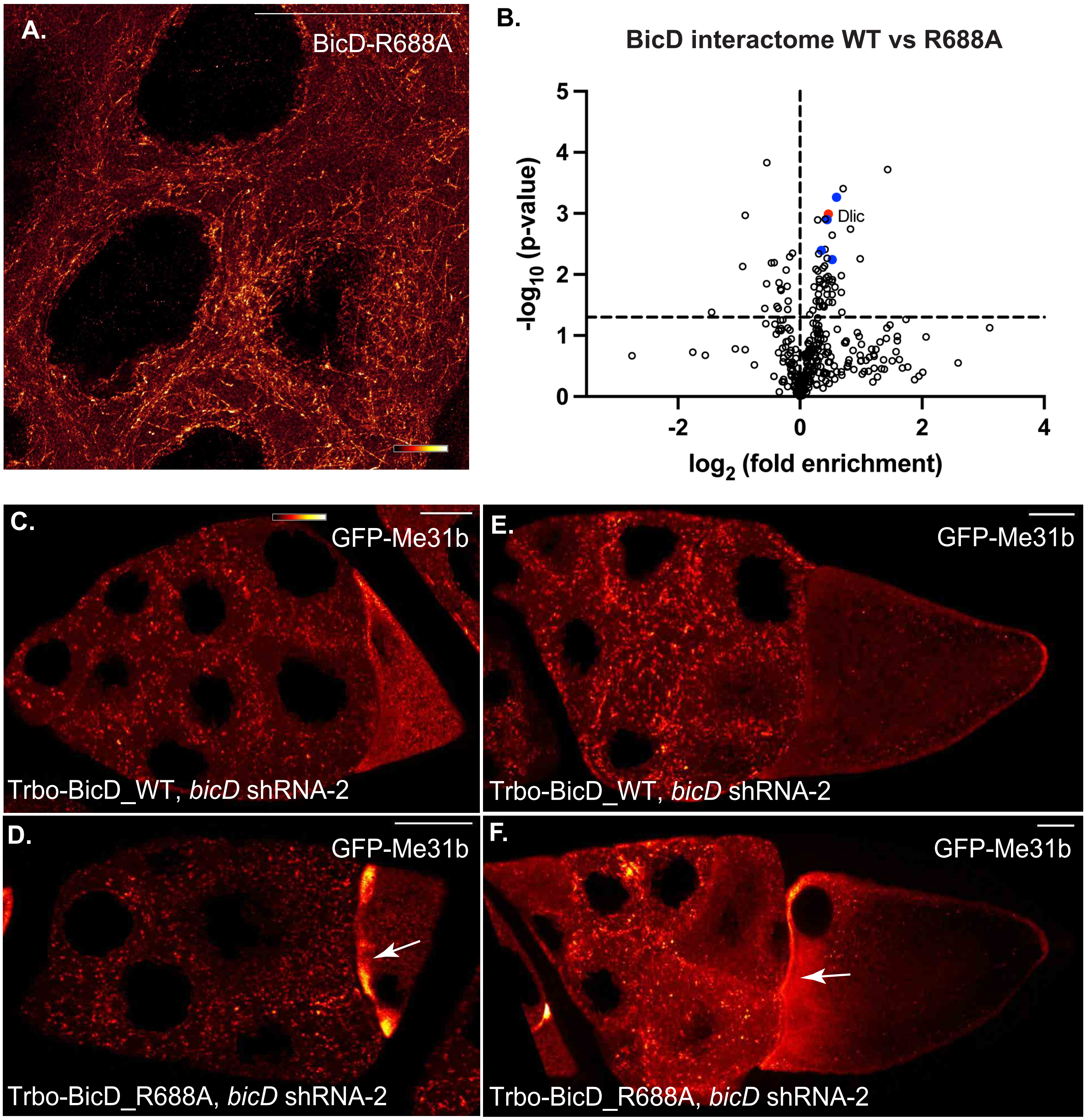


Supplemental Figure 3

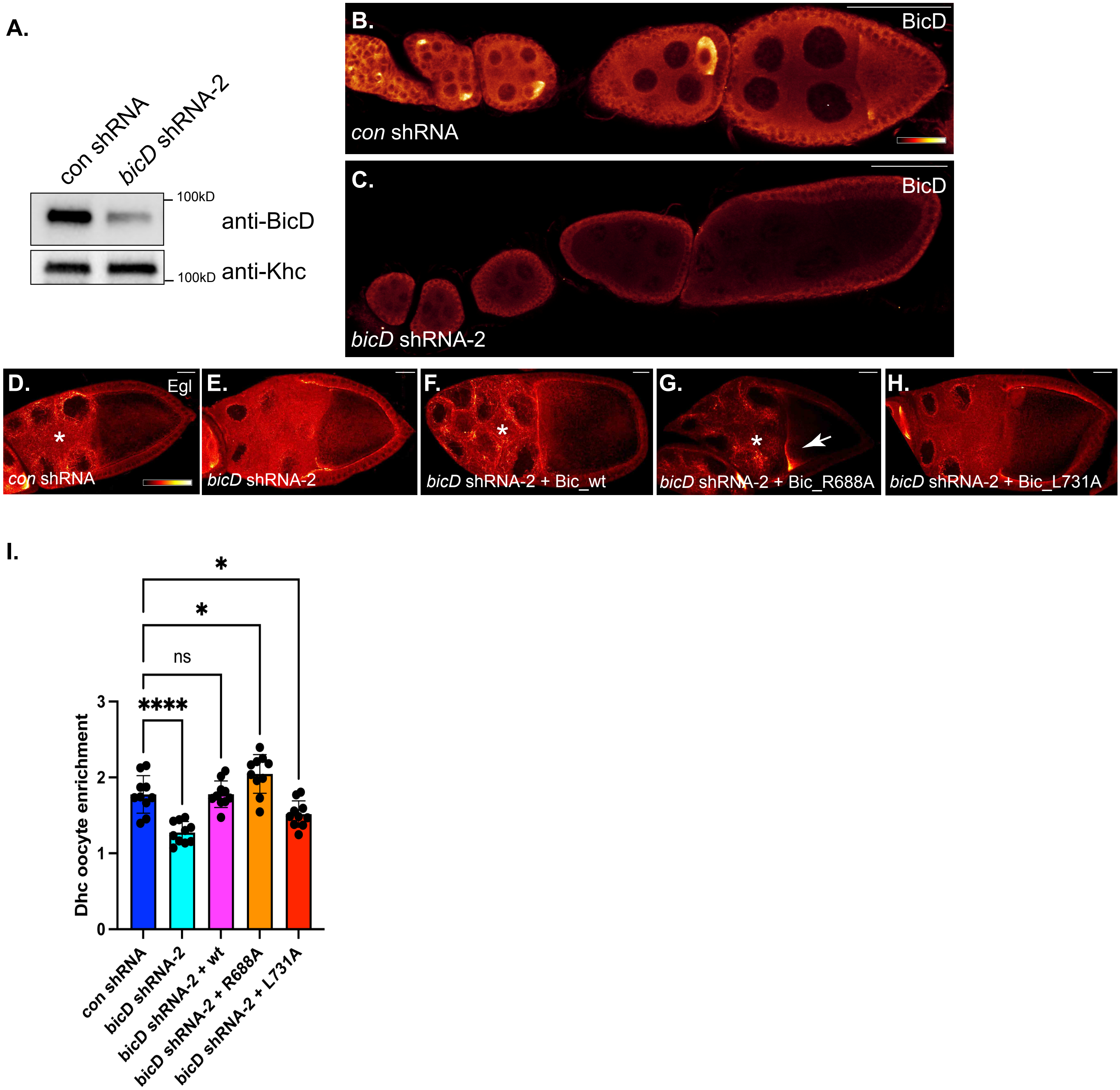


Supplemental Figure 4

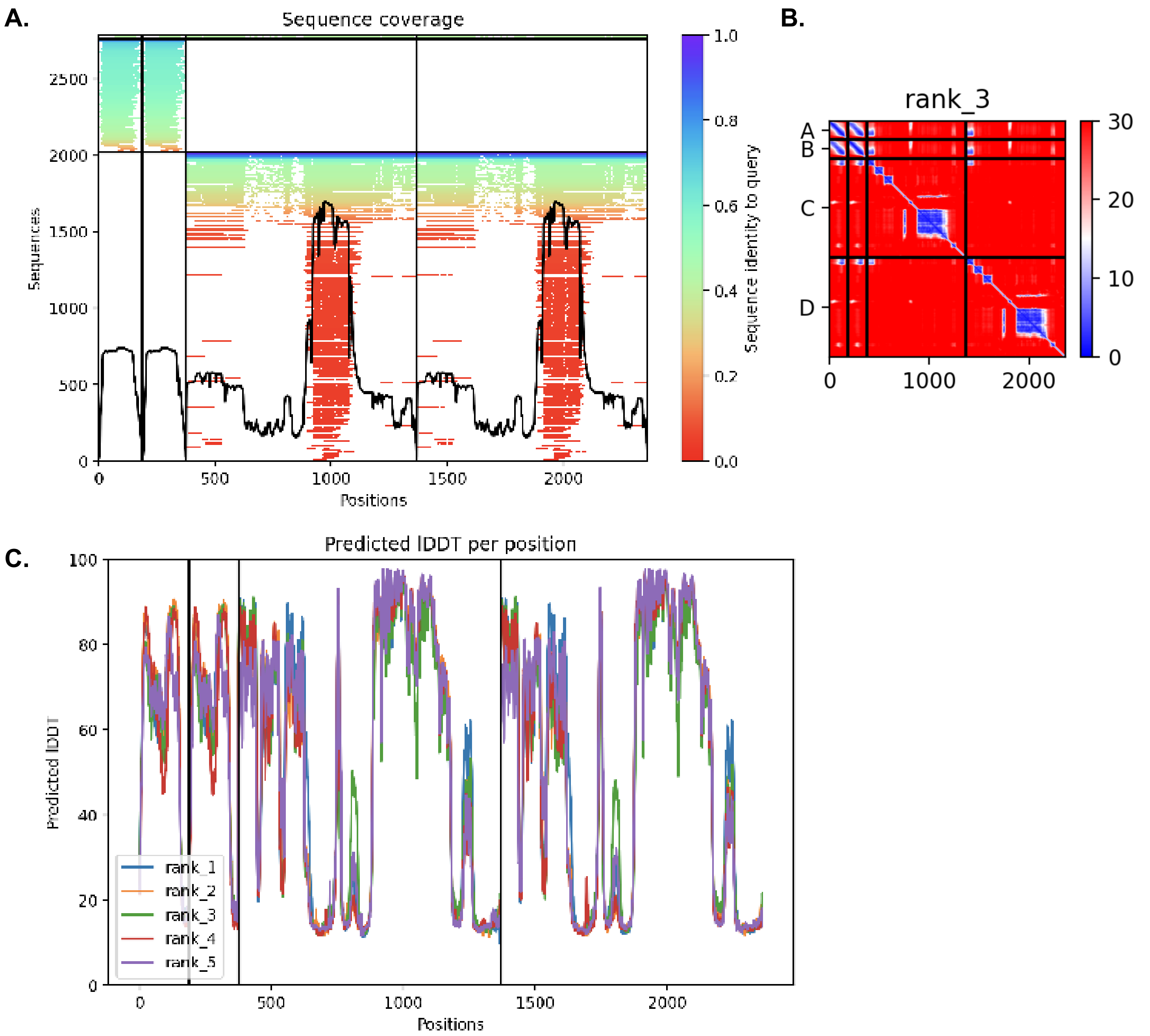


Supplemental Figure 5
